# New Data and Collaborations at the *Saccharomyces* Genome Database: Updated reference genome, alleles, and the Alliance of Genome Resources

**DOI:** 10.1101/2021.09.16.460706

**Authors:** Stacia R. Engel, Edith D. Wong, Robert S. Nash, Suzi Aleksander, Micheal Alexander, Eric Douglass, Kalpana Karra, Stuart R. Miyasato, Matt Simison, Marek S. Skrzypek, Shuai Weng, J. Michael Cherry

## Abstract

*Saccharomyces cerevisiae* is used to provide fundamental understanding of eukaryotic genetics, gene product function, and cellular biological processes. *Saccharomyces* Genome Database (SGD) has been supporting the yeast research community since 1993, serving as its de facto hub. Over the years, SGD has maintained the genetic nomenclature, chromosome maps, and functional annotation, and developed various tools and methods for analysis and curation of a variety of emerging data types. More recently, SGD and six other model organism focused knowledgebases have come together to create the Alliance of Genome Resources to develop sustainable genome information resources that promote and support the use of various model organisms to understand the genetic and genomic bases of human biology and disease. Here we describe recent activities at SGD, including the latest reference genome annotation update, the development of a curation system for mutant alleles, and new pages addressing homology across model organisms as well as the use of yeast to study human disease.

## INTRODUCTION

*Saccharomyces cerevisiae* is used to provide fundamental understanding of eukaryotic genetics, gene product function, and cellular biological processes. The published scientific literature from the yeast research community is integrated into the biomedical knowledgebase *Saccharomyces* Genome Database (SGD; www.yeastgenome.org). Biocurators with expertise in genetics, cell biology, and molecular biology have collected information from more than 100,000 published papers and combined the results from diverse experiments into a comprehensive resource for researchers, educators, and students.

The *S. cerevisiae* nomenclature has been maintained by SGD since 1993 (Cherry et al. 1997). Soon thereafter, SGD began providing the genetic and physical maps for the 16 yeast nuclear chromosomes (Cherry et al. 1997), the catalog of all known yeast proteins (Chervitz et al. 1999; Weng et al. 2003), biological process and molecular function annotations using the Gene Ontology (Dwight et al. 2001), as well as gene expression data and tools for analysis (Ball et al. 2001). SGD has maintained the reference genome from strain S288C, which was the first completely sequenced eukaryotic genome (Goffeau et al. 1996), and its annotation since 1998 (Cherry et al. 1998), along with sequence analysis and retrieval tools for studying that reference genome (Christie et al. 2004; Balakrishnan et al. 2005; Hirschman et al. 2006), and later broadened the reference panel by adding genomes of 11 additional highly studied strains to more fully support the work of the yeast research community (Engel and Cherry, 2013). In parallel to these activities, SGD also developed principles and practices for the extraction and curation of various types of biological data (Ball et al. 2000; Dwight et al. 2004; Hong et al. 2008; Engel et al. 2010; Costanzo et al. 2011; Park et al. 2012; Balakrishnan et al. 2013; Skrzypek and Nash 2015). SGD then turned to further enhancing existing tools and curation practices, including the development of an automated pipeline for pan-genome analysis (Song et al. 2015), release of the Variant Viewer for analysis of the reference genome panel (Sheppard et al. 2016), curation of the complete set of yeast metabolic pathways (Cherry 2015), and updated curation methods and data models for the capture of post-translational modifications (Hellerstedt et al. 2017), regulatory relationships (Engel et al. 2018), macromolecular complexes (Wong et al. 2019), and the yeast transcriptome (Ng et al. 2019).

As the de facto hub of the yeast research community, SGD also engages in a wide range of outreach and communication activities to disseminate published results, promote collaboration, facilitate scientific discovery, and inform users about new tools, data, or other database developments (MacPherson et al. 2017). These activities include participating in conferences and hosting workshops, direct contact with authors of yeast research papers, the posting of online help resources, and involvement in social media, including the production of video tutorials and webinars, all hosted on SGD’s YouTube channel (https://www.youtube.com/SaccharomycesGenomeDatabase). To increase readership and reach a broad audience, content posted on one outreach platform is often publicized or announced on other outreach platforms. We also collaborate with the *Genetics Society of America* to annotate online journal articles published in GENETICS and G3: Genes | Genomes | Genetics to associate yeast genes listed in these articles to their respective gene pages at SGD.

More recently, SGD and six other model organism-focused knowledgebases - Mouse Genome Database (MGD; http://www.informatics.jax.org, Bult et al. 2019), Rat Genome Database (RGD; http://rgd.mcw.edu, Laulederkind et al. 2018), Zebrafish Information Network (ZFIN; http://zfin.org, Ruzicka et al. 2019), WormBase (http://wormbase.org, Lee et al. 2018), FlyBase (http://flybase.org, Thurmond et al. 2019), and the Gene Ontology (GO) Consortium (http://www.geneontology.org, The Gene Ontology 2019) - have created the Alliance of Genome Resources (“the Alliance”; https://www.alliancegenome.org; Alliance 2019; Alliance 2022). Together, we are working to create an online resource that integrates, develops, and provides harmonized data for all member projects (Alliance 2020). The aim is to more broadly facilitate the use of model organisms to understand the genetic bases of human biology and disease. These efforts build on the well-established collaborations between these groups, who have long worked together to enhance data consolidation, dissemination, visualization, and the application of shared standards (i.e., “data harmonization”). Working within the Alliance has influenced SGD’s curation of mutant alleles and human diseases, as well as our use of homology throughout the SGD website to highlight the greater biological context of key findings in yeast research.

## GENOME VERSION R64.3.1

### Annotation updates and additions

The *S. cerevisiae* strain S288C reference genome annotation was updated for the first time since 2014 (Table 1). The new genome annotation is release R64.3.1, dated 2021-04-21. Resequencing of S288C in 2014 indicated that genomic sequence variation was less than expected between individual laboratory copies of this strain, illustrating that the underlying genome sequence is stable and complete. As such, while SGD updated the annotation of the genomic sequence, the fundamental sequence itself remains unchanged (Cherry et al. 1998; Engel et al. 2014). All new and updated annotations are sourced from the primary literature. The R64.3.1 update included the addition of seven open reading frames (ORFs), five noncoding RNAs (ncRNAs), two upstream ORFs (uORFs), and one long terminal repeat (LTR). Three ORFs had their translation starts shifted to a different methionine, and one ORF received a newly annotated intron and also had its translation stop shifted. We also changed the Sequence Ontology (SO; http://www.sequenceontology.org/; Eilbeck et al. 2005) term used to describe the non-transcribed spacers in the ribosomal DNA (rDNA) array.

**Table 1.** The *S. cerevisiae* strain S288C reference genome annotation was updated. The new genome annotation is release R64.3.1, dated 2021-04-21.

| Chromosome | Feature | Description of change | Reference |
| --- | --- | --- | --- |
| II | YNCB0008W aka GAL10-ncRNA | New ncRNA antisense to GAL10:<br>coordinates 276805..280645 | Houseley et al. 2008;<br>Pinskaya et al. 2009;<br>Geisler et al. 2012 |
| II | YNCB0014W aka TBRT/XUT_2F-154 | New ncRNA antisense to TAT1:<br>coordinates 376610..378633 | Awasthi et al. 2020 |
| III | RE/RE301 | New recombination enhancer:<br>coordinates 29108..29809 | Wu and Haber 1996 |
| V | YELWdelta27 | New Ty1 LTR:<br>coordinates 449274..449626 | Nene et al. 2018 |
| V | HPA3/YEL066W | Moved translation start to Met19:<br>old coordinates 26667..27206<br>new coordinates 26721..27206 | Sampath et al. 2013 |
| VII | OTO1/YGR227C-A | New ORF:<br>coordinates 949052..949225 Crick | Makanae et al. 2015 |
| VII | ROK1/YGL171W | Two new uORFs:<br>coordinates uORF1 182286..182407<br>coordinates uORF2 182291..182329 | Jeon and Kim 2010 |
| VIII | YHR052C-B | New ORF:<br>coordinates 212519..212692 Crick | He et al. 2018 |
| VIII | YHR054C-B | New ORF:<br>coordinates 214517..214690 Crick | He et al. 2018 |
| VIII | SUT169/YNCH0011W | New ncRNA:<br>coordinates 378254..379237 | Xu et al. 2009; Geisler<br>et al. 2012; Huber et<br>al. 2016; Bunina et al.<br>2017 |
| X | YJR012C | Moved start to Met76:<br>old coordinates 459795..460418 Crick<br>new coordinates 459795..460193 Crick | Sadhu et al. 2018 |
| X | YJR107C-A | New ORF<br>coordinates 628457..628693 Crick | Yagoub et al. 2015;<br>He et al. 2018 |
| XI | YKL104W-A | New ORF:<br>coordinates 245032..245286 | He et al. 2018 |
| XII | YLR379W-A | New ORF:<br>coordinates 877444..877716 | Internal reanalysis of<br>results from Song et<br>al. 2015 to find and<br>annotate missing<br>S288C ORFs |
| XII | NTS1-2, NTS2-1,<br>NTS2-2 | Change feature_type/SO_term from<br>SO:0001637 rRNA_gene to SO:0000183<br>non_transcribed_region |  |
| XIII | LDO45/YMR147W | Shift stop to be same as<br>LDO16/YMR148W, add intron:<br>old coordinates 559199..559870<br>new coordinates 559199..559780,<br>560156..560812 | Eisenberg-Bord et al.<br>2018 |
| XIII | YMR008C-A | New ORF:<br>coordinates 283081..283548 Crick | Internal reanalysis of<br>results from Song et<br>al. 2015 to find and<br>annotate missing<br>S288C ORFs |
| XIII | YNCM0001W aka<br>PHO84 lncRNA | New ncRNA:<br>coordinates 23564..26578 | Camblong et al. 2007 |
| XIV | LTO1/YNL260C | Move start to Met37:<br>old coordinates 156859..157455 Crick<br>new coordinates 156859..157347 Crick | Paul et al. 2015 |
| XVI | YNCP0002W aka<br>GAL4 lncRNA | New ncRNA:<br>coordinates 79562..82648 | Geisler et al. 2012 |

### New ncRNA systematic nomenclature

Also included in the R64.3.1 update is the establishment of a new systematic nomenclature for yeast ncRNAs (Table S1). For many years, a widely adopted systematic nomenclature has existed for yeast protein-coding genes, or ORFs, as many yeast researchers call them (Cherry et al. 1998). With this most recent genome version, we have established a systematic nomenclature for yeast ncRNAs that is similar to, but distinct from, that used for ORFs. All annotated *S. cerevisiae* ncRNAs are designated by a symbol consisting of four uppercase letters, a four-digit number, and another letter, as follows: Y for “Yeast”, NC for “noncoding”, A-Q for the chromosome on which the ncRNA gene resides (where “A” is chromosome I, “B” is chromosome II, etc., up to “P” for chromosome XVI, and lastly “Q” for the mitochondrial chromosome), a four-digit number corresponding to the sequential order of the ncRNA gene on the chromosome (starting from the left telomere and counting toward the right telomere), and W or C indicating whether the ncRNA gene is encoded on the “Watson” or “Crick” strand (where “Watson” runs 5′ to 3′ from left telomere to right telomere, and “Crick” runs 3’ to 5’). For example, <u>YNCP0002W</u> is the second ncRNA gene from the left end of chromosome XVI and is encoded on the Watson strand. Going forward, when evidence is published pointing to new ncRNA genes, they will be added to the annotation using the next sequential number available for the specific chromosome on which the ncRNA gene resides. In cases in which more than one ncRNA gene is added to any particular chromosome during the same annotation update (i.e., same genome revision), they will be named using the next sequential number starting with the leftmost ncRNA gene and proceeding to the right of the chromosome.

### Nomenclature updates - “legacy” gene names

SGD has long been the keeper of the official *Saccharomyces cerevisiae* gene nomenclature (Cherry 1998). Robert Mortimer handed over this responsibility to SGD in 1993 after maintaining the yeast genetic map and gene nomenclature for 30 years (Hawthorne and Mortimer 1960; Mortimer and Schild 1980). The accepted format for gene names in *S. cerevisiae* comprises three uppercase letters followed by a number. The letters typically signify a phrase (referred to as the “Name Description” in SGD) that provides information about a function, mutant phenotype, or process related to that gene, for example “ADE” for “ADEnine biosynthesis” or “CDC” for “Cell Division Cycle”. Gene names for many types of chromosomal features follow this basic format regardless of the type of feature named, whether an ORF, a tRNA, another type of non-coding RNA, an ARS, or a genetic locus. Some *S. cerevisiae* gene names that pre-date the current nomenclature standards do not conform to this format, such as *<u>MRLP38</u>, <u>RPL1A</u>*, and *<u>OM45</u>*. A few historical gene names predate both the nomenclature standards and the database, and are less computer-friendly than more recent gene names, due to the presence of punctuation. SGD recently updated these gene names to be consistent with current standards and to be more software-friendly by removing punctuation (Table 2).

**Table 2.** “Legacy” gene names which predate the database have been updated to be more software-friendly by removing unnecessary punctuation.

| ORF | Old gene name | New gene name |
| --- | --- | --- |
| YGL234W | ADE5,7 | ADE57 |
| YER069W | ARG5,6 | ARG56 |
| YBR208C | DUR1,2 | DUR12 |
| YIL154C | IMP2' | IMP21 |

Although non-standard historical names are maintained in SGD, any new names for yeast genes must conform to the standard format. The SGD Gene Registry (https://www.yeastgenome.org/reserved_name/new) is the agreed-upon system used by the *S. cerevisiae* community and SGD to reserve standard names for newly characterized genes. We have developed a set of Gene Naming guidelines and procedures in order to assist researchers in gene naming. We urge researchers to contact SGD prior to the publication of a new gene name (even if they have previously reserved it) to ensure that the gene name they wish to use is still appropriate. To date, 1363 currently annotated *S. cerevisiae* genes still have no standard gene name and are identified solely via their systematic names (ex. <u>YFL019C</u>). All *S. cerevisiae* ORFs are categorized into one of three groups: “verified” ORFs are those for which there is clear experimental evidence for the presence of expression of a protein-coding gene; “uncharacterized” ORFs are likely, but not yet established, to encode a protein; and “dubious” ORFs are unlikely to encode a protein (Fisk et al. 2006). Of the 1363 currently annotated *S. cerevisiae* ORFs without a standard gene name, 131 are verified, 571 are uncharacterized, and the remaining 661 are classified as dubious.

## ALLELES

To serve the needs of the yeast community more completely, and influenced by our active participation in the Alliance, SGD has recently revamped the way we curate mutant alleles of genes and has released new allele web pages. For over a decade, SGD has been recording mutant allele information as descriptive properties of phenotype annotations (Costanzo et al. 2009). We have recently elevated alleles in the database from phenotype properties to primary data objects, *i.e*., standalone entities to which other annotations, such as phenotypes or genetic interactions, can be attached, and that have their own detailed web pages.

Previously, as part of phenotype curation, only allele names, mutant types, and free-text descriptions were collected. We now have expanded the information gathered and displayed for alleles to include affected gene, alias names, description in free text including relevant nucleotide and/or amino acid changes and correspondence to human alleles when known, and allele type. Allele type, which is captured using the *structural_variant* (SO:0001537) branch of the Sequence Ontology, describes the change to the relevant sequence feature with terms like *missense_variant* (SO:0001583), *stop_gained* (SO:0001587), and *frameshift_variant* (SO:0001589). To further describe the functional effect of the structural change within the context of phenotype annotations, SGD uses the *functional_effect_variant* (SO:0001536) branch of the SO, which contains terms like *null_mutation* (SO:0002055), *loss_of_function_variant* (SO:0002054), and *dominant_negative_variant* (SO:0002052).

### New allele pages

The new allele pages at SGD contain different types of information divided into several sections: Overview, Phenotypes, Genetic Interactions, Shared Alleles network, external Resources, and relevant Literature (fig. 1). The Allele Overview provides general information about the allele, including its name, the affected gene, the type of allele (*e.g*., *missense_variant*, as described above), and a description of sequence change and/or domain mutated. The Phenotype Annotations for an allele include an observable feature (*e.g*., “cell shape”), a qualifier (*e.g*., “abnormal”), a mutant type (e.g., null), strain background, and a reference. In addition, annotations are classified as classical genetics or high-throughput (*e.g*., large scale survey, systematic mutation set). The Genetic Interactions for an allele are defined as experimentally observed genetic interactions between that allele and another of a different gene. All interactions listed in SGD are curated by BioGRID (Oughtred et al. 2021). The Shared Alleles network displays positive genetic interactions, negative genetic interactions, and phenotypes that are shared between the given allele and other alleles (fig. 1). The Resources section provides links to allele-related information, such as mutant strain repositories, external phenotype and interaction databases, and information about the Yeast Phenotype Ontology. Additionally, all literature associated with an allele can be found on its Literature page (fig. 1).

**Fig. 1.**
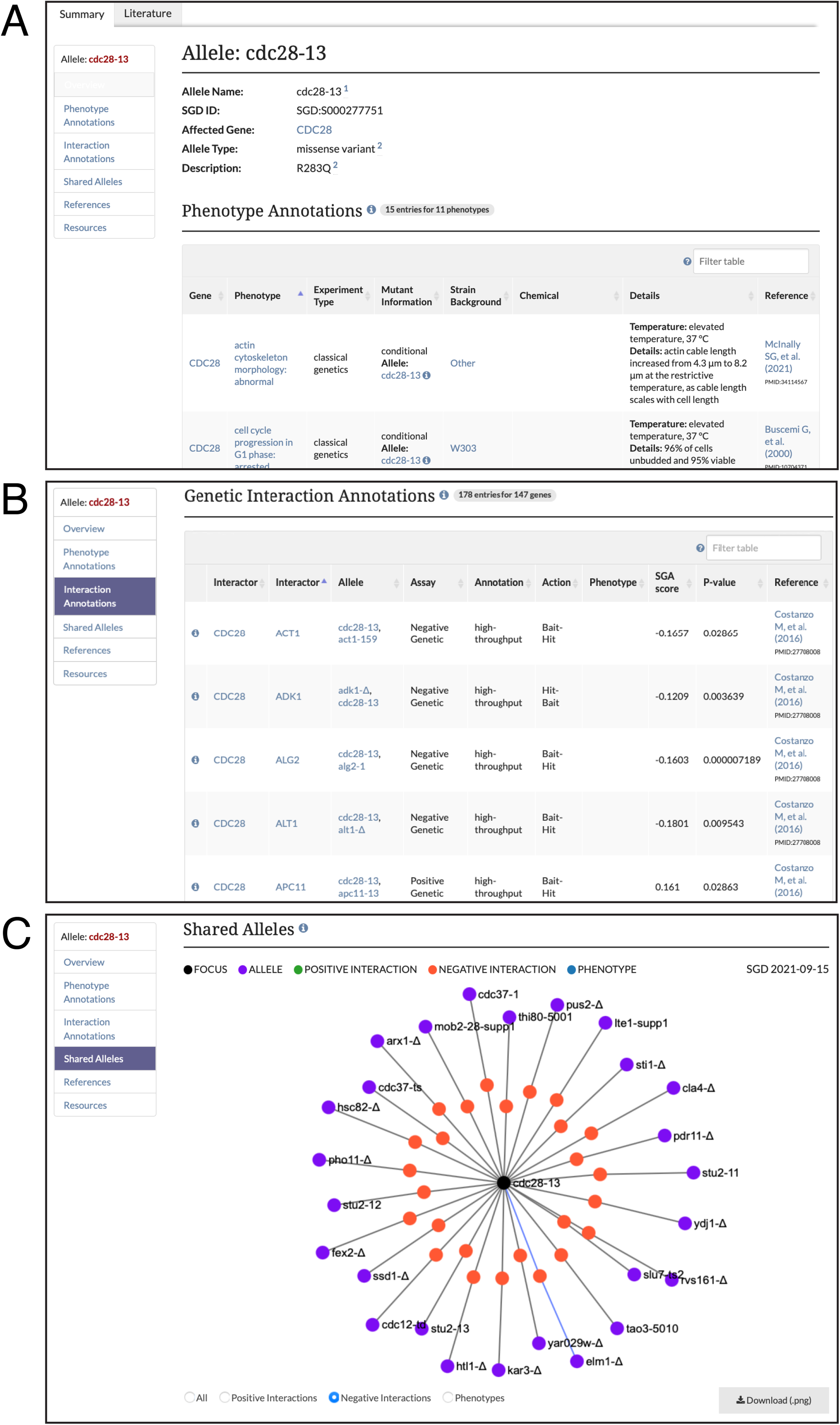
SGD allele pages include (A) Overview with name, affected gene, type of allele, and a description of sequence change and/or domain mutated; phenotype annotations; (B) genetic interaction annotations; and (C) Shared Alleles network diagram depicting shared phenotypes and interactions with other alleles.

### Updates to Interactions data/pages

To further accommodate the enhanced focus on alleles and for improved clarity, SGD has made recent updates to the way we present Interaction data. Previously, genetic and physical interaction annotations were combined in one table. These annotations are now displayed in separate annotation tables on Interactions pages (ex. https://www.yeastgenome.org/locus/S000003424/interaction), Reference pages (ex. https://www.yeastgenome.org/reference/S000305076), and in YeastMine (https://yeastmine.yeastgenome.org/yeastmine/templates.do) to allow for the listing of allele designations, Synthetic Genetic Analysis (SGA) Scores, and P-values from the global genetic interactions paper by Costanzo et al. (2016).

### Searching for alleles

All alleles for a specific gene can be accessed via the Alleles section on the Locus Summary Page or downloaded from YeastMine using the Genes -> Alleles template (https://yeastmine.yeastgenome.org/yeastmine/template.do?name=Gene_Alleles&scope=all). The alleles have also been added to SGD’s Elasticsearch, with ‘facets’ for publications, allele types, affected genes, and phenotypes, which allow for the browsing, partitioning, and viewing of the approximately 14,000 yeast alleles annotated so far in SGD.

## HOMOLOGY AND THE ALLIANCE OF GENOME RESOURCES

SGD and six other model organism focused knowledgebases - MGD, RGD, ZFIN, WormBase, FlyBase, and the GO Consortium - have recently created a new knowledgebase for model organisms. These major community-driven model organism projects have taken up the mission of harmonizing common data types and creating an integrated web resource. The Alliance of Genome Resources has brought together biocurators and software engineers from all the MODs to build a new central resource. Teams have been created to define data models for the major data types provided by the MODs. These teams also define how the information should be presented. Software engineers from the MODs work together to create the cloud computational environment, component tools, and web pages that define such an important resource. Gene products, proteins, ncRNAs, and pseudogenes are connected by their homology, molecular functions, biological processes, cellular component location, anatomical expression, and association with disease. Human gene details are provided by RGD.

One of the first work products to come out of the Alliance was a consolidated set of orthologs, using data from several different computational and manually curated sources (Howe et al. 2018). Many aspects of data integration presented at the Alliance require a common set of orthology relationships among genes for the organisms represented, including human. The Alliance provides the results of all methods that have been benchmarked by the Quest for Orthologs Consortium (QfO; https://questfororthologs.org/, Linard et al. 2021). The homolog inferences from the different methods have been integrated using the DRSC Integrative Ortholog Prediction Tool (DIOPT; Hu et al. 2011), which integrates a number of existing methods including those used by the Alliance: Ensembl Compara (Vilella et al. 2009), HUGO Gene Nomenclature Committee (HGNC; Povey et al. 2001), Hieranoid (Kaduk and Sonnhammer 2017), InParanoid (O’Brien et al. 2005), the Orthologous MAtrix project (OMA; Schneider et al. 2007), OrthoMCL (Li et al. 2003), OrthoFinder (Emms and Kelly 2015), OrthoInspector (Linard et al. 2011), PANTHER (Mi et al. 2018), PhylomeDB (Huerta-Cepas et al. 2008), Roundup (Deluca et al. 2006), TreeFam (Li et al. 2006), and ZFIN (Westerfield et al. 1997). This set of assertions (the “orthology set”) is key because it allows the inference of function from one species to another, and is especially helpful because work using a highly studied and experimentally tractable organism such as yeast can be informative regarding the biology of other organisms in which targeted experiments are not possible, such as human (Dolinski and Botstein 2007). Currently included in the Alliance and in the orthology set are five other leading model organisms in addition to *S. cerevisiae* (yeast): *Caenorhabditis elegans* (worm), *Drosophila melanogaster* (fruit fly), *Danio rerio* (zebrafish), *Rattus norvegicus* (rat), and *Mus musculus* (mouse).

SGD takes advantage of the homology data from the Alliance to provide easy access to information about homologous genes in just one click. At SGD, we have used the orthology set to provide links between SGD gene pages and those for orthologous genes at the Alliance (fig. 2). On SGD gene pages, users will find hexagonal icons representing each model organism (human, mouse, rat, zebrafish, fly, worm, and yeast) for which there is homologous gene information at the Alliance. Clicking on the icon immediately directs the user to the gene page for the selected model organism on the Alliance website. The Alliance gene pages present a variety of data types for all of the various organisms, including functional annotations using the Gene Ontology, phenotypes, disease associations, alleles, variants, sequence features, expression information, and both physical and genetic interactions. Data for individual genes can be downloaded directly from Alliance gene pages, and bulk downloads are available from the Alliance Downloads site (https://www.alliancegenome.org/downloads).

**Fig. 2.**
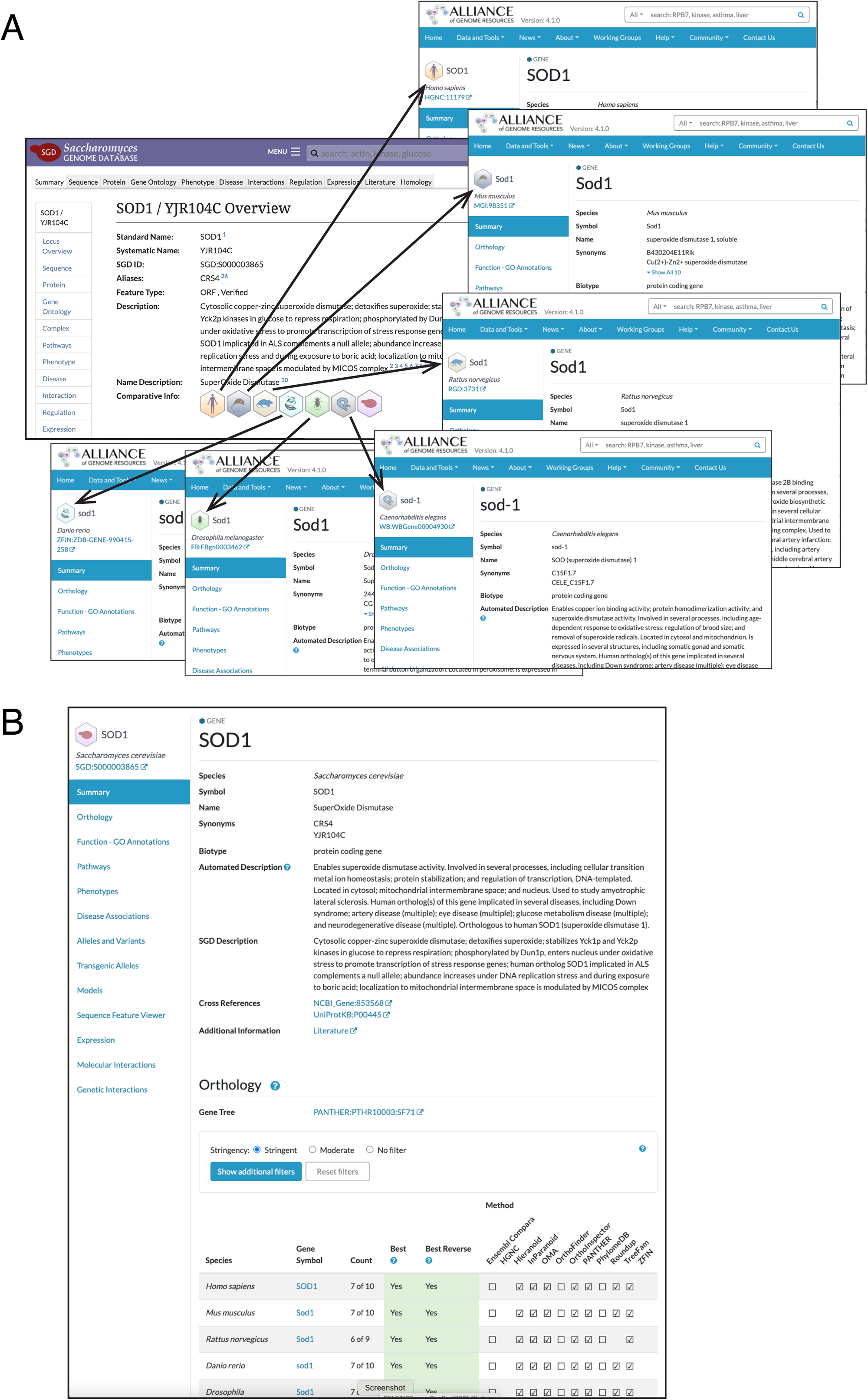
(A) SGD provides links between SGD gene pages and those for orthologous genes at the Alliance of Genome Resources. Hexagonal icons represent each organism (human, mouse, rat, zebrafish, fly, worm, and yeast) for which there is gene information at the Alliance. (B) Curated information from SGD is presented on yeast gene pages at the Alliance of Genome Resources. The various types of yeast data that can be found at the Alliance can be viewed using the menu on the left side of the gene page, and include orthologs, functional annotations using the Gene Ontology, cellular pathways, phenotypes, disease associations, alleles, variants, sequence features, expression, and interactions. Data can be downloaded in bulk from https://www.alliancegenome.org/downloads.

### Homology pages

We have also used the Alliance orthology set to establish new Homology pages at SGD for protein-coding genes. The information displayed on the Homology Pages is divided into several sections: Homologs, Functional Complementation, Fungal Homologs, and External Identifiers (fig. 3). The Homologs section lists information about genes in the Alliance orthology set, including species, gene ID, and gene name, with links to the corresponding gene page at the Alliance. The Functional Complementation section presents data about cross-species complementation between yeast and other species, as curated by SGD and including legacy data from the Princeton Protein Orthology Database (P-POD; Heinicke et al. 2007). These curated Functional Complementation data are also displayed on SGD Reference pages. The Fungal Homologs section shows curated information including species, gene ID, and database source for orthologs in 24 additional species of fungi, such as those in *Candida, Neurospora*, and *Aspergillus*, among others. The External Identifiers section lists identifiers for the protein from various database sources.

**Fig. 3.**
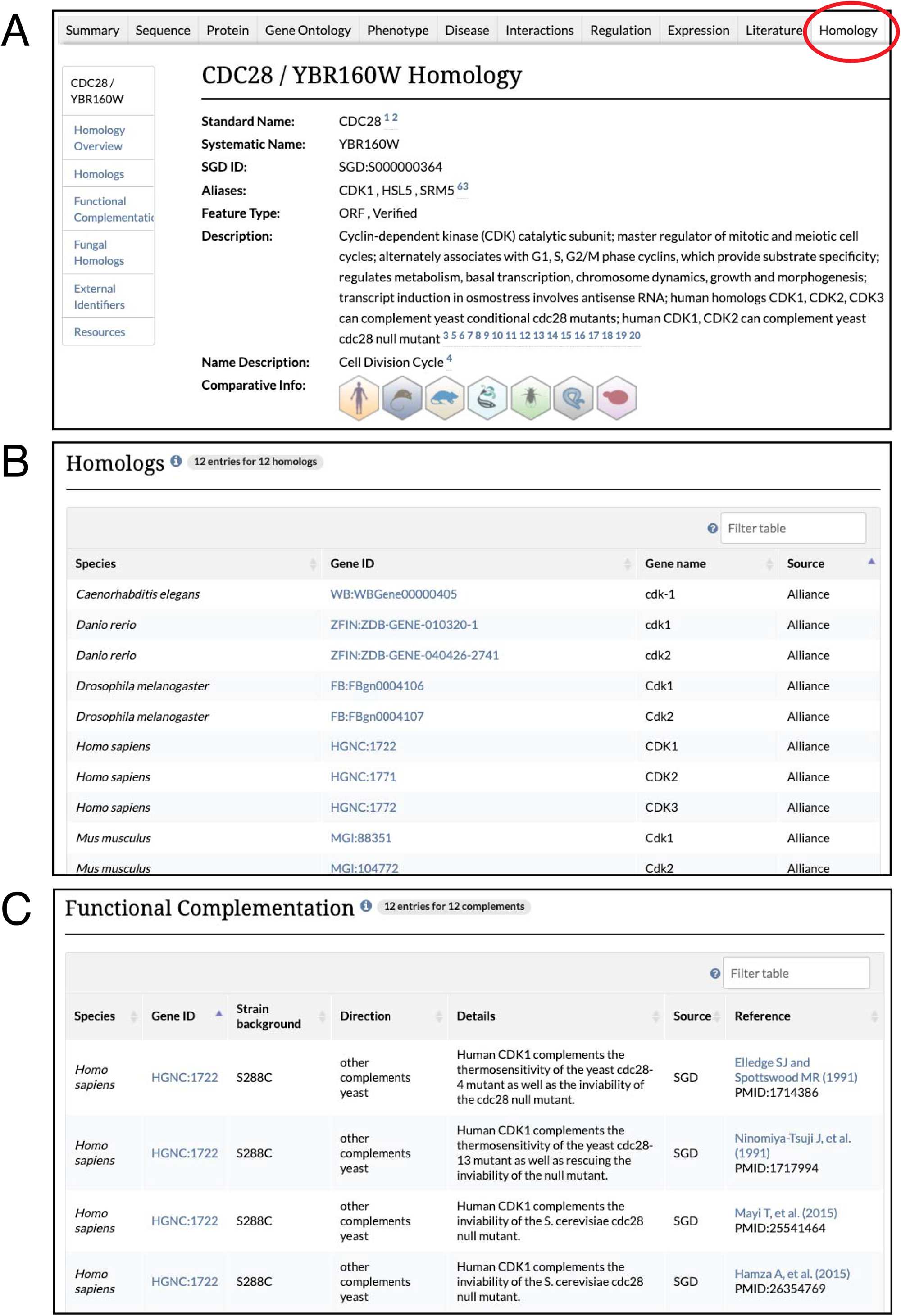
SGD homology pages (A) include an Overview of general information about the yeast gene, with links to homologous gene pages at the Alliance; (B) information about known homologs including species corresponding Gene ID, and name of the homolog; and (C) data about cross-species functional complementation between yeast and other species. Also included on the page, but not shown here, are Fungal Homologs, gene identifiers in other databases, and links to external resources.

### Disease pages

To promote the use of yeast as a catalyst for biomedical research, SGD utilizes the Disease Ontology (DO; Schriml et al. 2018) to describe human diseases that are associated with yeast homologs. Disease Ontology annotations to yeast genes are now available through SGD’s new Disease pages, each of which corresponds to a Disease Ontology term, such as *amyotrophic lateral sclerosis* (DOID:332), and lists out all yeast genes annotated to the term by SGD. Yeast genes with one or more human disease associations also have a new Disease tab accessible from the genes’ respective locus pages (fig. 4). The Disease tab shows all manually curated, high-throughput, and computational disease annotations for the yeast gene. Additionally, these pages feature a network diagram that depicts shared disease annotations for other yeast genes and their human homologs.

**Fig. 4.**
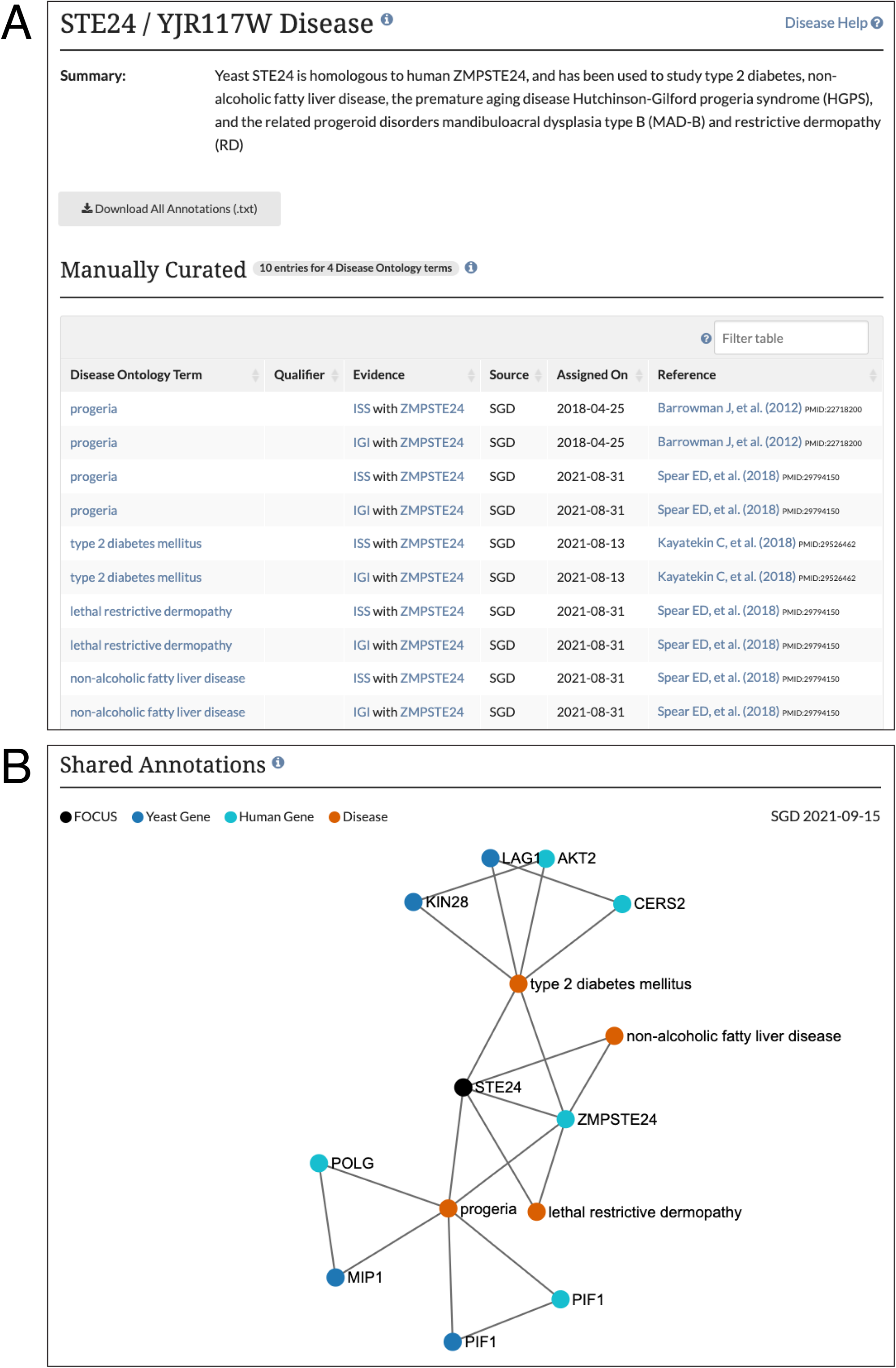
SGD disease pages use the Disease Ontology to describe human diseases that are associated with yeast homologs, and include (A) disease annotation summary and individual annotations; and (B) shared annotation network that depicts shared disease annotations for other yeast genes and their human homologs.

While the Alliance will provide much of the information researchers will want to explore, there will continue to be data that cannot be harmonized across organisms because of its unique characteristics, such as an aspect of biology only found in one of the species or the nuance of an experimental assay that is unique to one community. In time, these unique bits of information will be provided directly from the Alliance site. However, until then, the main knowledgebases are essential. All of the MODs look forward to the common user interface and ease of discovering common associations between genes in other organisms. This is an important step to allow researchers, educators, and students to have access to the gold standard of expertly curated information on each of these foundations of biological science.

## OTHER UPDATES

### Supplemental data and published datasets on reference pages

SGD procures and displays supplemental materials for references stored in our database. We are hosting data from past, present, and future papers on our literature pages. To access these data, simply search SGD with the PubMed ID and then look for the “Downloadable Files.” Additionally, published large-scale datasets from Gene Expression Omnibus (GEO) are displayed in the “Published Datasets” section. The dataset title is linked to a dataset-specific page and a controlled vocabulary of terms is used to bin similar datasets into broad categories.

### Submit data form

Authors can submit data and information about their publications, including novel results, datasets (including accession IDs), or other important information, using SGD’s simple “Submit Data” form (https://www.yeastgenome.org/submitData).

### Explore button

SGD introduced a new “Explore SGD” button on our homepage, which allows users to peruse SGD data and pages without an initial search query. After selecting the “Explore SGD” button, users are redirected to a search results page where they can browse all of the information in SGD. The tool is designed for both new and veteran users alike, as new users are provided a glimpse into the warehouse of information SGD contains, while seasoned users may discover something new. After clicking on the “Explore SGD” button, use the categories on the left to navigate through the various pages and examine areas of interest. An “Explore” link has also been added to the selection of links available in the black bar at the top of every SGD webpage, giving users the ability to access the search results page from anywhere on the SGD website.

### microPublication Biology

SGD has partnered with *microPublication Biology* (https://www.micropublication.org) to promote the growing model of rapid publication of research results. *microPublication* further expands this rapid-release model by coupling publication with curation and deposition into community-directed authoritative databases such as SGD and the other Alliance member databases mentioned above. Each publication is peer-reviewed, assigned a digital object identifier (DOI), published online, indexed in PubMed (https://pubmed.ncbi.nlm.nih.gov), and then integrated into the relevant databases, speeding information dissemination and scientific discovery. SGD encourages authors to utilize *microPublication* for new research findings or reagents, experimental results that do not fit into a larger existing or future narrative, negative results, successful replications of recent work, cautionary findings regarding unsuccessful attempts at replication of recent work, “data not shown” from other publications, and data from student projects. All published yeast articles are immediately available in SGD (https://www.yeastgenome.org/search?category=reference&journal=microPublication.%20Biology).

## ACKNOWLEDGEMENTS

Thanks to Jodi Lew-Smith for help preparing this manuscript.

